# Improving power while controlling the false discovery rate when only a subset of peptides are relevant

**DOI:** 10.1101/2020.10.20.347278

**Authors:** Andy Lin, Deanna L. Plubell, Uri Keich, William S. Noble

## Abstract

The standard proteomics database search strategy involves searching spectra against a peptide database and estimating the false discovery rate (FDR) of the resulting set of peptide-spectrum matches. One assumption of this protocol is that all the peptides in the database are relevant to the hypothesis being investigated. However, in settings where researchers are interested in a subset of peptides, alternative search and FDR control strategies are needed. Recently, two methods were proposed to address this problem: subset-search and all-sub. We show that both methods fail to control the FDR. For subset-search, this failure is due to the presence of “neighbor” peptides, which are defined as irrelevant peptides with a similar precursor mass and fragmentation spectrum as a relevant peptide. Not considering neighbors compromises the FDR estimate because a spectrum generated by an irrelevant peptide can incorrectly match well to a relevant peptide. Therefore, we have developed a new method, “filter then subset-neighbor search” (FSNS), that accounts for neighbor peptides. We show evidence that FSNS properly controls the FDR when neighbors are present and that FSNS outperforms group-FDR, the only other method able to control the FDR relative to a subset of relevant peptides.

## 1 Introduction

In a typical proteomics database search, mass spectra are searched against a database consisting of peptides reasonably expected to be found in the sample. For example, mass spectra generated from human cells would be searched against the human proteome. Following the database search, target-decoy competition is used to estimate the false discovery rate (FDR) of a set of peptide-spectrum matches (PSMs).^1,2^ By setting an FDR threshold, typically 1%, we can identify a set of confidently detected PSMs. Although this process is standard across the field and is valid for many proteomic analyses, there are situations where this FDR control strategy can be problematic.

Specifically, the standard process implicitly assumes that all peptides in a sample are relevant to a given hypothesis. This assumption does not hold when researchers are only interested in a subset of the peptides present in the sample. For example, scientists may only be interested in a single protein, a single type of post-translational modification, a single pathway, or a single organism in a microbial community. In such settings, variants of the FDR control process are needed to ensure proper FDR control while trying to maximize the number of discoveries among the relevant subset of peptides.

We consider a peptide to be *relevant* when the detection, or lack of detection, of that peptide is pertinent to the hypothesis being asked. On the other hand, irrelevant peptides are peptides that are inconsequential to the question being asked. A common class of irrelevant peptides is human contaminants, typically keratin, which can be artificially introduced into a non-human sample during sample preparation. Human keratin contaminants are irrelevant because detecting these peptides does not affect the biological interpretation of the data. In this scenario, the peptide database will mostly consist of relevant peptides, since the list of contaminants is typically small.

However, there are other scenarios where the proportion of relevant peptides in the protein database is small. One example is when proteomics is used to study the biology of *Plasmodium falciparum*, the causative agent of malaria. When *P. falciparum* is studied in a laboratory, it must be cultured in a medium containing human red blood cells. Therefore, any *P. falciparum* sample will also contain human peptides. In this setting, the detection of human peptides is irrelevant since the goal of the experiment is to study the biology of *P. falciparum*. In practice, researchers search their experimental spectra against the concatenated human and *P. falciparum* proteomes. Since the human proteome is much larger than the *P. falciparum* proteome, the proportion of relevant peptides in the combined database is small.

For this work we focus on the setting where the proportion of relevant peptides in the database is exceedingly small. Generally this would occur when investigators are pursuing a focused biological hypothesis such as the effect of a drug on a single molecular pathway or the effect of a perturbation on a single organism in a microbial community. One concrete example is the detection of the protein toxin ricin, RCA60, for law enforcement purposes. Detection of the ricin toxin is important because the possession, transfer, or use of this toxin is federally regulated throughout most of the world. The challenge for law enforcement is that the castor plant and the seeds of the castor plant, where ricin is expressed, are not regulated. In fact, there are many legitimate reasons to possess and transfer the plant and seeds. For example, the castor plant is a common ornamental plant, and castor seeds are used in the production of castor oil. As a result, prosecutors must be able to directly detect the ricin toxin. All other proteins expressed in the castor plant are not useful for prosecution. This means that a single protein is relevant while the entire remaining proteome is irrelevant.

One very commonly used, yet incorrect, strategy found in the literature, which we call “search-then-select,” involves estimating FDR on PSMs from the full database but only reporting the relevant PSM subset.^3,4^ Search-then-select does not properly control the FDR because the relevant PSM score distribution does not necessarily match the irrelevant PSM score distribution.^5,6^ For example, in the case where researchers are interested in post-translationally modified peptides the score distribution of irrelevant unmodified peptides can differ from relevant modified peptides.^7,8^

Motivated by this problem, three methods have been proposed for calculating the statistical significance of a subset of relevant PSMs (Table 1 and Figure 1). One method is to simply search the spectra against the database consisting only of relevant peptides (“subset-search”).^9,10^ A second method searches the spectra against relevant and irrelevant peptides but applies FDR control only to relevant PSMs (“group-FDR”^11^) (also sometimes referred to as “separate FDR”^5^). Finally, in the third method, spectra are searched against all peptides but FDR is controlled with respect to only the set of relevant peptides using information from all PSMs (“all-sub”)^5,8,12^ For this analysis, we chose the method from Sticker *et al*. as a representative example of this type of method.^12^

**Figure 1:**
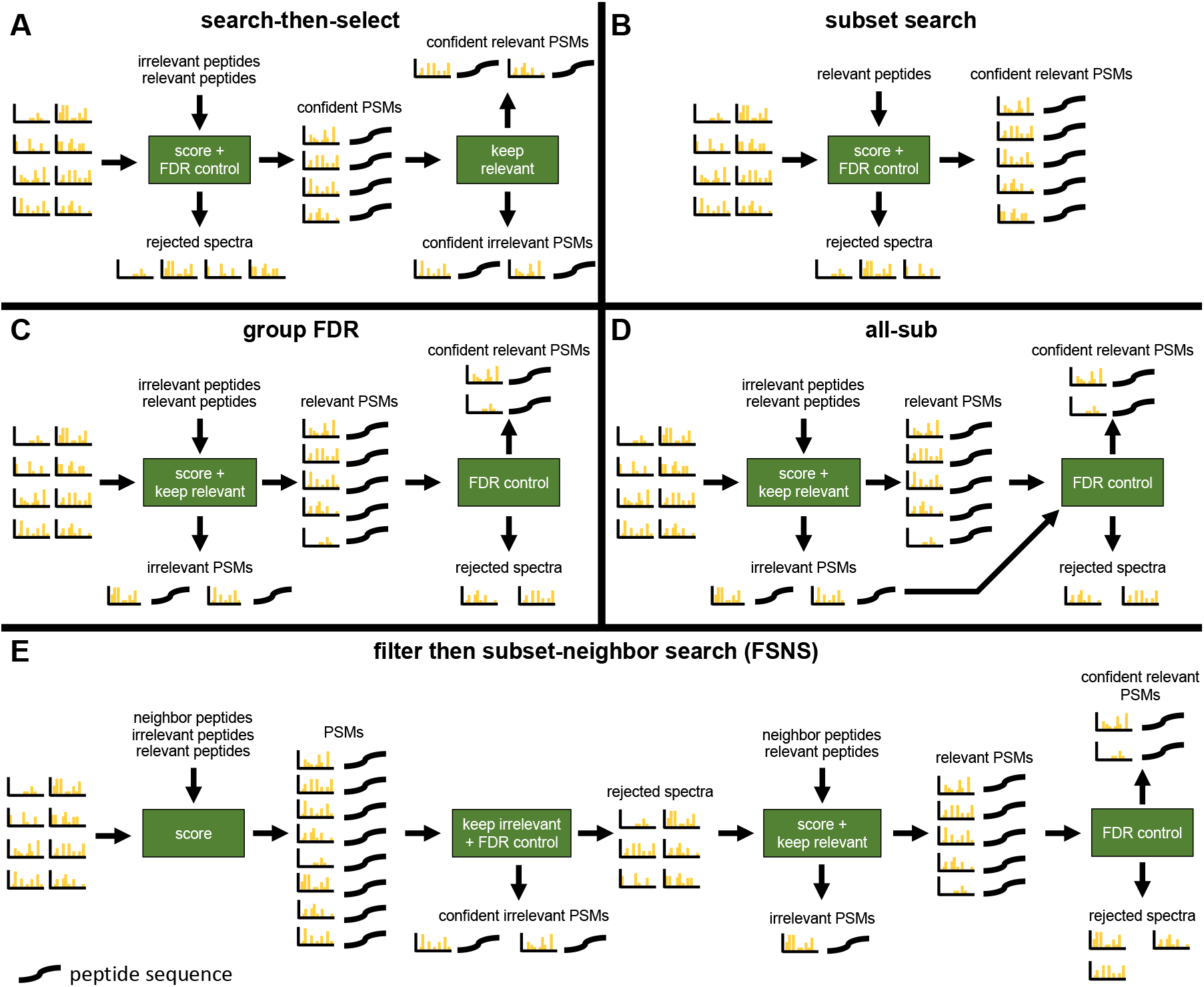
Graphical overview of methods. “Keep relevant” means that any PSM that involves a relevant peptide is kept while every other PSM is removed. “Keep irrelevant” means any PSM that involves an irrelevant peptide is kept while every other PSM is removed. Neighbors peptides are defined and explicitly considered separate from irrelevant peptides only in “filter then subset-neighbor search” (FSNS).

**Table 1:** Summary of methods. We represent irrelevant peptides with an ‘I’, neighbor peptides with an ‘N’, and relevant peptides with an ‘R’. If a database consists of multiple groups of peptides, then a ‘+’ is used. Therefore, a database consisting of both relevant and neighbor peptides would be represented as ‘R+N’. Group-FDR is with respect to R for the “Database to Search” step.

|  | search-then-select | subset-search | all-sub | group-FDR | FSNS |
| --- | --- | --- | --- | --- | --- |
| Consider neighbor peptides? |  |  |  |  | ✓ |
| Initial filtering step? |  |  |  |  | R+I+N |
| Database to search | R+I+N | R | R+I+N | R+I+N | R+N |
| Apply group-FDR? |  |  | variation | ✓ | ✓ |

Our interest in this problem was rekindled when analyzing data where the only relevant peptides were those of the ricin protein. This is an example where the relevant peptides comprise a very small subset of the peptides in the database. However, with this data we also encountered a new phenomenon—”neighbor” peptides—that has not previously been explicitly taken into account in this context. A neighbor peptide is an irrelevant peptide that has a similar precursor mass and fragmentation (MS2) spectrum as a relevant peptide. As explained below, ignoring the existence of neighbors peptides can compromise an FDR controlling procedure.

Our investigation of existing analysis procedures starts by evaluating their ability to control the FDR. Others have already pointed out that search-then-select fails to control the FDR.^5–8^ Here we also demonstrate that all-sub can fail to properly control the FDR when the relevant peptides comprise a small subset of the peptides in the database. In contrast, our initial analysis indicated that group-FDR and subset-search do not fail this test. However, subsequent analysis shows that subset-search begins to fail when a sufficient number of neighbor peptides are thrown into the mix. Specifically, in the analysis of our ricin data, we observe cases where a spectrum generated by a neighbor of a ricin peptide offers a very good match for the corresponding ricin peptide, even though the latter may not be in the sample. These potentially incorrect PSMs cannot be accounted for by the target-decoy competition in subset-search, since this process does not account for the existence of peptides that are not relevant but are neighbors.

These experiments left us with group-FDR as the only procedure that properly controls the FDR for the general case of searching a small subset of relevant peptides. However, like search-then-select and all-sub, group-FDR suffers from the problem of searching the relevant spectra against a large irrelevant database, thereby compromising its power.10 This observation motivated our introduction of a new method, called “filter then subset-neighbor search” (FSNS), that tries to retain most of subset-search’s ability to limit the search to the relevant part of the database while fixing the latter’s failure to control the FDR by explicitly accounting for neighbor peptides (Table 1 and Figure 1).

Our analysis shows that FSNS offers greater statistical power than group FDR, i.e., FSNS typically delivers more discoveries than group FDR at a fixed FDR threshold. Given that none of the other methods offer robust FDR control, FSNS is our recommended method when a small subset of the peptides in a sample are relevant.

## 2 Methods

Notation is summarized in Table 2.

**Table 2:**
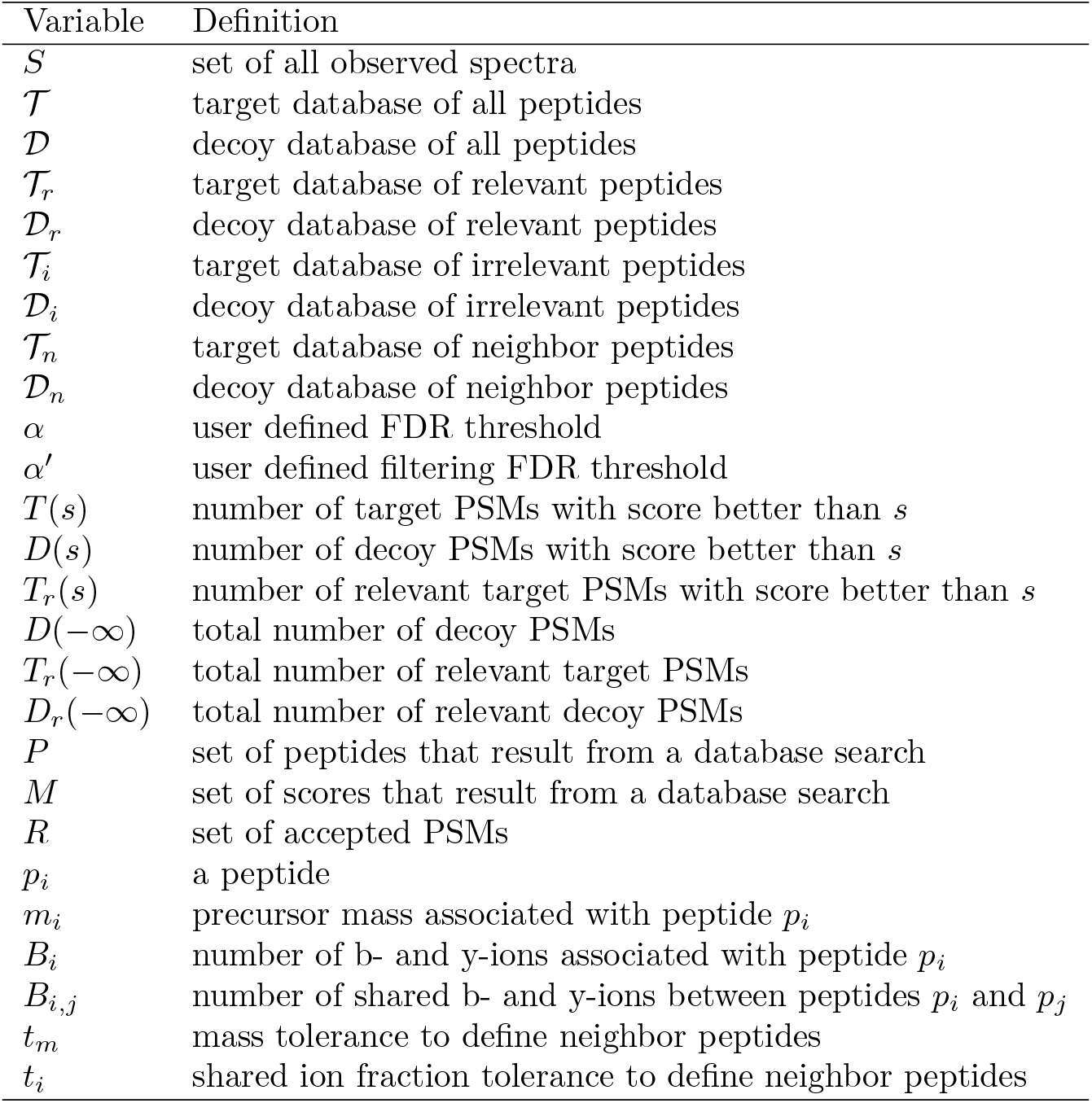
Variables and their definitions. Note that there is no overlap between the relevant and irrelevant peptides. Thus, 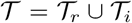 and 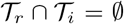. In addition, if neighbor peptides are not defined, then they are considered to be irrelevant. However, if neighbor peptides are defined, then they are considered distinct from irrelevant peptides.

| Variable | Definition |
| --- | --- |
| $S$ | set of all observed spectra |
| $\mathcal{T}$ | target database of all peptides |
| $\mathcal{D}$ | decoy database of all peptides |
| $\mathcal{T}_r$ | target database of relevant peptides |
| $\mathcal{D}_r$ | decoy database of relevant peptides |
| $\mathcal{T}_i$ | target database of irrelevant peptides |
| $\mathcal{D}_i$ | decoy database of irrelevant peptides |
| $\mathcal{T}_n$ | target database of neighbor peptides |
| $\mathcal{D}_n$ | decoy database of neighbor peptides |
| $\alpha$ | user defined FDR threshold |
| $\alpha'$ | user defined filtering FDR threshold |
| $T(s)$ | number of target PSMs with score better than $s$ |
| $D(s)$ | number of decoy PSMs with score better than $s$ |
| $T_r(s)$ | number of relevant target PSMs with score better than $s$ |
| $D(-\infty)$ | total number of decoy PSMs |
| $T_r(-\infty)$ | total number of relevant target PSMs |
| $D_r(-\infty)$ | total number of relevant decoy PSMs |
| $P$ | set of peptides that result from a database search |
| $M$ | set of scores that result from a database search |
| $R$ | set of accepted PSMs |
| $p_i$ | a peptide |
| $m_i$ | precursor mass associated with peptide $p_i$ |
| $B_i$ | number of b- and y-ions associated with peptide $p_i$ |
| $B_{i,j}$ | number of shared b- and y-ions between peptides $p_i$ and $p_j$ |
| $t_m$ | mass tolerance to define neighbor peptides |
| $t_i$ | shared ion fraction tolerance to define neighbor peptides |

### 2.1 Neighbor peptides

We say that two peptides are “neighbors” if their masses are similar and if they share sufficiently many singly-charged b- and y-ions. More formally, we say peptides *p*_1_ and *p*_2_ are neighbors if (1) the difference in the associated peptide masses *m*_1_ and *m*_2_, specified in units of ppm, is less than a specified mass tolerance

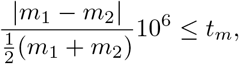

and (2) the proportion of b- and y-ions shared by *p*_1_ and *p*_2_ is greater than a specified fraction *t_i_*:

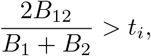

where *B*_1_ (respectively, *B*_2_) is the number of possible singly-charged b- and y-ions that can be associated with peptide *p*_1_ (*p*_2_), and *B*_12_ is the number of shared such ions between the two peptides. Here, two (theoretical) ions are considered shared if their m/z values, discretized to bins of size 0.05 Da, fall in the same bin. In this work, we set *t_m_* equal to twice the precursor mass tolerance used in the associated database search, and we set *t_i_* = 0.25.

### 2.2 FDR control methods

We consider five methods for controlling the FDR (the first four of which have been previously published, Table 1 and Figure 1). Pseudocode descriptions of each algorithm can be found in the supplement.

#### Search-then-select

This method takes as input a set *S* of observed spectra, a database 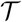 of target peptides, a corresponding database 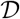 of decoy peptides, a database 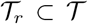 of relevant peptides, and a confidence threshold *α*. The spectra are searched against the concatenated target-decoy database 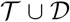, yielding an optimal PSM for each spectrum. The FDR for the set of PSMs is calculated using

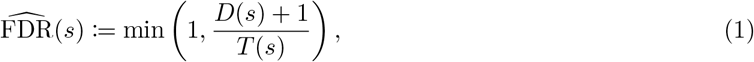

where *T*(*s*) is the number of target PSMs with score ≥ *s* and *D*(*s*) is the number of decoy PSMs with score ≥ *s*. Note that the min operation is used to prevent the estimated FDR from exceeding 100%. In the algorithm, the set of confident discoveries *A* = *A*(*α*) is defined as all PSMs whose score is ≥ *s_α_*, where

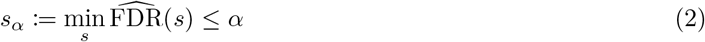

and *α* is the user defined FDR threshold (typically 1%). Thus far, the procedure corresponds to the standard target-decoy competition protocol for FDR control.^1,2^ In the search-then-select protocol, the subset of target peptides in *A* that do not involve a relevant peptide are filtered out, leaving only target PSMs involving peptides of interest. This final set of PSMs is designated as the set *R* of accepted PSMs.

#### subset-search

The subset-search method^9,10^ is similar to the standard target-decoy competition protocol, except that it operates only on the set of relevant peptides 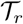. The procedure takes as input the observed spectra *S*, a set of relevant target peptides 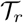, the set of corresponding decoy peptides 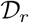, and the significance threshold *α*. The FDR is estimated using (1), and the discovery list *A* is determined using the score cutoff *s_α_* from (2).

#### Group-FDR

In the group-FDR method^11^ (also known as separate FDR^5^), the inputs include observed spectra *S*, a relevant target database 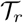, a relevant decoy database 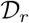, an irrelevant target database 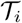, an irrelevant decoy database 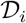, and a threshold *α*. In this search, neighbor peptides 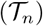 are absorbed into the irrelevant peptide set 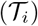; therefore 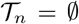. A database search is conducted against a concatenated database consisting of all targets and decoys 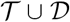 (which is the same as 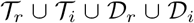). After the search, PSMs that involve irrelevant peptides or their associated decoys are filtered out. The FDR is then estimated on the remaining set of PSMs using (1), and the discovery list is determined using the score cutoff *s_α_* defined by (2).

#### All-sub

The all-sub method12 is conceptually similar to group-FDR except for a variation in how the FDR is estimated. In particular, the input to all-sub is the same as for group-FDR. Spectra are searched against the concatenated target-decoy database 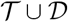, but instead of using (1), the FDR is estimated using

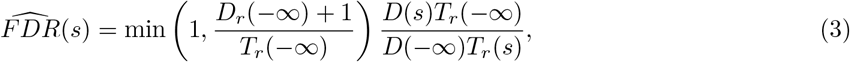

where *T_r_*(*s*) is the number of relevant target PSMs with scores ≥ *s, T_r_*(-∞) is the total number of relevant target PSMs, *D_r_*(−∞) is the total number of relevant decoy PSMs, and D(-∞) is the total number of decoy PSMs.

#### Filter then subset-neighbor search (FSNS)

As its name suggests, FSNS is the only method presented here that explicitly accounts for neighbors. The input for FSNS includes, in addition to the input to group-FDR 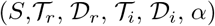, a “filtering” FDR threshold *α′*, and an optionally non-empty database of neighbor peptides 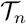 together with their corresponding decoys 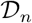.

FSNS employs an initial filtering step in which the spectra are searched against all the peptides: 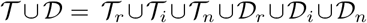. Using (1), FSNS then estimates the FDR among the PSMs involving only peptides in 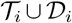. Subsequently, all spectra that are identified at the filtering threshold of *α′* are then removed from further consideration because they are deemed to be most likely generated by irrelevant peptides. For this study we set the filtering FDR threshold *α′* to be 0.01.

All remaining spectra are then searched using the group FDR protocol, with the neighbors playing the role of the irrelevant peptides. Specifically, all remaining spectra are searched again, this time against a database of relevant and neighbor peptides 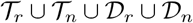. Following the second database search, PSMs that involve neighbor peptides or their associated decoys are removed from consideration, and the list of discoveries is determined from (1) and (2), as described above, using threshold α.

### 2.3 Datasets

To evaluate the various FDR control methods, we use four tandem mass spectrometry datasets (Table 3, Supplemental Table 1-3), which have been deposited in the PRIDE Archive (http://www.ebi.ac.uk/pride/archive) with the dataset identifier XXXXX.^1^

**Table 3:** UPS1 and yeast data. The table list the number of scans found in each UPS1 and yeast run. In the database search, the relevant file on the left was concatenated to the irrelevant file on the right. Note that all file names start with “UWPRLumos_20190515_DP_DDA_”.

| relevant file | scans | irrelevant file | scans |
| --- | --- | --- | --- |
| UPS_1_22.ms2 | 10552 | yeast_1_45.ms2 | 71631 |
| UPS_2_23.ms2 | 10178 | yeast_2_46.ms2 | 71611 |
| UPS_3_24.ms2 | 9794 | yeast_3_47.ms2 | 71931 |

#### UPS1/Yeast

This dataset consists of six mass spectrometry runs. Three runs came from a yeast whole cell lysate, and the remaining three runs came from the Universal Proteomics Standard Set 1 (UPS1, Sigma-Aldritch). The resulting spectra were then merged to create three *in silico* mixtures.

To prepare the yeast sample, yeast strain BY4741 (MATa his3Δ1 leu2Δ0 met15Δ0 ura3Δ0) (Dharmacon) was cultured in YEPD to mid-log phase, harvested, and lysed with 8M urea buffer solution and bead beating (7 cycles of 4 minutes beating with 1 min rest on ice). The resulting cell lysate was reduced, alkylated, and digested for 16 hours. Next, the peptide digest was desalted using a mixed-mode (MCX) method, dried down overnight via speedvac, and brought up with synthetic iRT peptide standards (Pierce Peptide Retention Time Calibration Mixture) to 1*μ*g/*μ*l total proteome. A bicinchoninic acid assay (Pierce BCA Protein Assay Kit) was used to determine total protein content.

To prepare the UPS1 sample, the Universal Proteomics Standard Set 1 (Thermo Scientific) was reduced, alkylated, and digested for 16 hours in the same manner as the yeast sample.

For both prepared samples, peptides were separated with a Waters NanoAcquity UPLC and emitted into a Orbitrap Fusion Lumos (Thermo Scientific, San Jose, California). Pulled tip columns were created from 75 *μ*m inner diameter fused silica capillary (New Objectives, Woburn, MA) in-house using a laser pulling device and packed with 3 *μ*m ReproSil-Pur C18 beads (Dr. Maisch GmbH, Ammerbuch, Germany) to 30 cm. Trap columns were created from 150 *μ*m inner diameter fused silica capillary fritted with Kasil on one end and packed with the same C18 beads to 3 cm. Solvent A was 0.1% formic acid in water and solvent B was 0.1% formic acid in 98% acetonitrile. For each injection, 3 *μ*l (approximately 1 μg total protein on column) was loaded and eluted using a 90-minute gradient from 5 to 35% B, followed by a 40 minute wash. Data were acquired using data-dependent acquisition (DDA).

To acquire DDA data, the Orbitrap Fusion Lumos was set to positive mode in a top-20 configuration. Precursor spectra 400–1600 m/z) were collected at 60,000 resolution to hit an AGC target of 3 × 10^6^. The maximum inject time was set to 100 ms. Fragment spectra were collected at 15000 resolution to hit an AGC target of 10^5^ with a maximum inject time of 25 ms. The isolation width was set to 1.6 m/z with a normalized collision energy of 27. Only precursors charged between +2 and +4 that achieved a minimum AGC of 5 × 10^3^ were acquired. Dynamic exclusion was set to “auto” and to exclude all isotopes in a cluster. Both the UPS1 peptides and the yeast peptides were injected into the mass spectrometer three times (Table 1).

#### UPS1/Yeast

A second UPS1/yeast dataset was used for a power analysis, and the data for this dataset was downloaded from PRIDE (project number PXD001819).14 In this study, UPS1 proteins were spiked into a yeast cell lysate at nine different concentrations: 50 amol/*μ*g, 125 amol/*μ*g, 250 amol/*μ*g, 500 amol/*μ*g, 2.5 fmol/*μ*g, 5 fmol/*μ*g, 12.5 fmol/*μ*g, 25 fmol/*μ*g, 50 fmol/*μ*g. Three technical replicates were generated for each of the nine samples. Runs corresponding to UPS1 spike-in concentrations of ≤2.5 fmol/*μ*g were removed from consideration because these runs had zero confident UPS1 peptide detections at 1% FDR. This resulted in 12 usable runs for our analysis.

The yeast lysate was created in an 8M urea/0.1 M ammonium bicarbonate buffer at a protein concentration of 8 *μ*g/*μ*L. UPS1 proteins were spiked into 20 *μ*g of the yeast lysate to create nine different concentrations of UPS1. Following the spike-in step, the sample was reduced and alkylated. Digestion was done overnight using trypsin in a 1M urea buffer. Following the digestion step, each sample was desalted and analyzed in triplicate on a nanoRS UHPLC system (Dionex, Amsterdam, The Netherlands) coupled to an LTQ-Orbitrap Velos mass spectrometer (Thermo Fisher Scientific, Bremen, Germany) in top 20 data-dependent acquisition mode with dynamic exclusion set to 60 seconds.

Two *μ*L of each sample were loaded into a C-18 column (75 *μ*m IDx15 cm, in-house packed with C-18 Reprosil) where solvent A consists of 5% acetonitrile and 0.2% formic acid and solvent B consists of 80% acetonitrile and 0.2% formic acid. The flow rate was 300 nL/min flow rate. For the first 75 minutes of the gradient the percentage of solvent B increased from 5% to 25%. During the next 30 minutes, solvent B increased to 50% and finally during the last 10 minutes solvent B increased to 100%. MS1 scans were acquired in the Orbitrap on the 300–2000 m/z range with the resolution set to 60,000.

#### Ricin

Data for this dataset was downloaded from PRIDE (project ID PXD007933). In this study, castor seeds from various castor plant cultivars were collected for sample processing.15 Crude castor seed extracts were prepared from castor seeds via five different methods, designated M0 through M4. Method M0 involves mashing the seed to a semi-uniform consistency. M1 first removes the seed coat by soaking the seeds in sodium hydroxide, then mashes the peeled seeds. M2 takes the product from M1 and washes it with acetone. M3 and M4 involve a protein precipitation step where either magnesium sulfate or acetone, respectively, is used. Following preparation of the crude castor seed extracts, the extracts were inactivated by heating to 100 C for 1 hour.

The crude castor seed extracts were placed in a buffer (PBS from 10x concentrate, Fluka, containing 11.9 mM phosphates, 137 mM sodium chloride, 2.7 mM potassium chloride, pH 7.4, to which 0.01% Tween-20 was added) and then centrifuged for 10 minutes at 4°C and 16,000 g. Following centrifugation, the resulting aqueous layer was incubated with urea and 500 mM dithiothreitol (DTT) for 60 minutes at 37°C. Then, 400 mM iodoacetamide (IAA) was added and then incubated in the dark at 37°C for 60 m to alkylate cysteine thiol groups. Once alkylated, samples were diluted using ammonium bicarbonate buffer to which calcium choloride was added. Samples were digested overnight at 30°C using trypsin. After digestion, samples were acidified and desalted by solid phase extraction.

A Waters NanoAcquity dual pump LC system (Waters, Milford, Massachusetts) was used to separate peptides on the column. Peptides were separated on a fused silica capillary column which consisted of a trapping column (4 cm x 150 *μ*m inner diameter) and an analytical column (70 cm x 75 *μ*m inner diameter, 360 *μ*m outer diameter). Peptides were eluted at a rate of 300 nL/min for 150 m using a nonlinear gradient. A Q Exactive Plus or Q Exactive HF mass spectrometer (Thermo Scientific, San Jose, California) was used to collect the data. Both MS and MS/MS spectra was collected at high resolution. Specifically, the Q Exactive Plus precursor spectra were acquired at 35,000 resolution (mass resolving power) for precursor spectra and MS/MS spectra at 17,500 resolution, while on the Q Exactive HF, the resolution settings were 60,000 for precursor spectra and 15,000 for MS/MS spectra.

The ricin dataset consists of 54 different runs of castor seed extracts, corresponding to multiple replicates for each of the five sample preparation protocols, and are summarized in Supplemental Table 2.

#### Human

A subset of 12 human mass spectrometry runs from PRIDE project PXD011189 were downloaded. This study developed a new sample preparation protocol, sample preparation by easy extraction and digestion (SPEED),16 and compared it to three previously developed methods of filter-aided sample preparation (FASP), single-pot solid-phase-enhanced sample preparation (SP3), and urea-based in-solution digestion (Urea-ISD). To generate biomass HeLA cells (ATCC CCL-2) were grown in DMEM media at 37 °C and supplemented with 10% FCS and 2 mm l-Glutamine. After sample collection, cells were washed with PBS and then pelleted and stored at −80 °C until the lysis step.

In the SPEED method, samples were resuspended in trifluoroacetic acid (TFA) and then incubated at room temperature for 2 minutes. After the samples were neutralized using a solution of 2 M TribaseNext and TFA, Tris(2-carboxyethyl)phosphine (TCEP) and 2-Chloroacetamide (CAA) was added. After the TCEP and CAA was added the samples were incubated for 5 min at 95 °C. Following the incubation step, a trypsin digestion step was conducted for 20 hour at 37 °C. Finally, the peptide mixture was acidified and desalted.

In FASP,^17^ samples were suspended in a solution of 4% SDS, 100 mm Tris/HCl, and 100 mm DTT, incubated at 95 °C for 5 min, and then sonicated at 4 °C to induce lysis. Following the lysis step, samples were processed using a Microcon-30kDa Centrifugal Filter Units (Merck). With this unit the sample underwent several rounds of centrifugation in a solution of 8 M urea and 0.1 M Tris-HCl. During one round of centrifugation the peptide were alkylated using IAA. Following all rounds of centrifugation, the filter device was rinsed and desalted. The filtrate were digested for 20 h at 37 °C using trypsin at a protein/enzyme ratio of 50:1.

In the SP3 method,^18^ cells were lysed in a solution of 1% SDS, 1x complete Protease Inhibitor Mixture (Roche, Basel, Switzerland), and 50 mm HEPES buffer. During the lysis step the samples were incubated at 95 °C for 5 min and further sonicated for 300 seconds at 4 °C. Following the lysis step samples were reduced and alkylated using DTT and IAA, respectively. Then, 2 **μ*L* of paramagnetic beads was added to the mixture. After the beads were immobilized by a magnetic rack for two minutes, they were washed and resuspended in a trypsin solution and digestion was carried out for 20 h at 37 °C. Peptides were eluted off the beads using a 2% dimethyl sulfoxide in water.

In the urea-ISD method, cells were lysed by suspending the sample in a pH 8 solution of 8 m urea, 50 mm Tris-HCl, and 5 mm DTT. Then, it was sonicated for 10 min at 4 °C. Following the lysis step, the samples were incubated for 1 h at 37 °C and then centrifuged for 5 min. Samples were alkylated for 30 min at room temperature in the dark using IAA. Proteins were digested using trpysin for 20 hours at 37 °C in a urea solution with 50 mm Tris-HCl.

Once all the samples had been prepared they were all analyzed on a EASY-nanoLC 1200 (Thermo Fisher Scientific) coupled online to a Q Exactive Plus mass spectrometer (Thermo Fisher Scientific). One *mu* g of sample was injected into a 50 cm Acclai PepMap column (Thermo Fisher Scientific) using a linear 180 min gradient of 3 to 28% acetonitrile in 0.1% formic acid at a 200 nL/min flow rate. The Q Exactive Plus was operated in a top 10 data-dependent acquisition mode with a dynamic exclusion window of 30 seconds and collected scans in the m/z range of 300–1650. MS1 scans were acquired with a resolution of 70,000 and fragment scans were recorded with a resolution of 17,500.

For all datasets, files in .ms2 format^19^ were generated from the Thermo RAW file vendor format using Proteowizard version 3.0.^20^

### 2.4 Database search

For each database search, two protein databases were employed, one designated “relevant” and one “irrelevant” (Table 4). The databases for the castor plant (which contains the ricin protein), yeast strain ATCC 204508, and human were downloaded from Uniprot (https://www.uniprot.org/) in March 2018, May 2019, and January 2019, respectively. Protein sequences for the UPS1 proteins were downloaded from the Sigma Aldrich website (https://www.sigmaaldrich.com) in December 2018. The UPS1 and yeast sequences were concatenated into a single protein database. Each protein database was digested to peptides *in silico* using the tide-index tool in Crux version 3.2, allowing up to one missed cleavage, up to three methionine oxidations, and with clipped N-terminal methionines.^21,22^ For the UPS1/yeast database we considered UPS1 peptides to be relevant and yeast peptides irrelevant. For the castor plant database we considered the ricin protein relevant and all other castor proteins irrelevant. Finally, for the human database we considered a single randomly chosen detectable human protein as relevant and all other human proteins irrelevant. This process was repeated five times for the human database (Uniprot ID: P18206, Q9Y490, P07900, P08238, and P10809). For the UPS1/yeast and human database, peptides found in both relevant and irrelevant proteins were removed from the analysis. On the other hand, castor plant peptides found in both relevant (ricin) and irrelevant (non-ricin) proteins were considered relevant. In all cases, decoy peptides were generated by randomly shuffling the target sequence while keeping the N-terminal and C-terminal amino acids fixed. This was done to maintain the amino acid frequencies of the N-terminus, C-terminus, and internal amino acids. A different decoy database was generated for each run, yielding 54 decoy castor plant databases, three decoy UPS1/yeast databases for the concatenated runs, 12 decoy UPS1/yeast databases for the previously published runs, and 60 decoy human databases.

**Table 4:** Databases used in database searches. For the UPS1/yeast database and the five iterations of the human database, peptides in common between the relevant and irrelevant database were removed from analysis. Any relevant ricin peptide also found in the non-ricin castor plant proteome was considered to be relevant.

| relevant database | peptides | irrelevant database | peptides |
| --- | --- | --- | --- |
| UPS1 | 2,552 | yeast | 611,861 |
| ricin | 80 | non-ricin castor plant | 2,340,370 |
| human protein P18206 | 364 | all other human proteins | 2,653,948 |
| human protein Q9Y490 | 506 | all other human proteins | 2,653,757 |
| human protein P07900 | 102 | all other human proteins | 2,654,127 |
| human protein P08238 | 90 | all other human proteins | 2,654,130 |
| human protein P10809 | 136 | all other human proteins | 2,654,122 |

Database searches were conducted using Tide with the combined p-value score function23 against a concatenated target-decoy database. The precursor mass tolerances, estimated by Param-Medic,24 were found to be 31 ppm for the concatenated UPS1/yeast runs, 80 ppm for the castor plant runs, 63 ppm for the previously published UPS1/yeast runs, and 38 ppm for the human runs. All Tide parameters were set to their default values, except an isotope error of 1 was allowed and the “top-match” parameter (i.e., number of reported PSMs per spectrum) was set to 10,000. A post-processing script (Supplemental File 1) implemented each FDR estimation method. Once the list of PSMs was finalized, the Crux assign-confidence command was used to estimate FDR for all methods except for all-sub. Each PSM is assigned a q-value where the value is the minimum FDR at which the PSM is confidently detected. For all-sub, we created our own Python implementation of this method to estimate all-sub q-values (Supplemental File 2).

### 2.5 Evaluating the validity of FDR control methods

To gauge whether each of the considered methods properly controls the FDR we tested whether the empirical mean of the false discovery proportion (FDP) is significantly different than the FDR threshold. The idea is that since the FDR is defined as the expectation of the FDP, controlling the FDR at level *α* implies that the empirical mean of the FDP, averaged across multiple independent runs, should converge to a number ≤ *α*. In particular, that empirical mean should not exceed α in a statistically significant manner.

Here we computed the empirical mean of the FDP by randomly dividing our data into ten, roughly equal parts, and we used a heuristic explained below to reliably approximate the FDP in each of the ten runs and hence its mean over the same ten runs.

We then performed a one-sided t-test asking whether the observed mean of the FDP is significantly larger than α. If the answer was positive then we had a reason to doubt the validity of the proposed FDR-controlling method. By the same token, an insignificant deviation does not prove the validity of the method, but it does lend some confidence in it.

To simulate multiple independent runs we randomly split a UPS1 and yeast run into 10 equal parts. After splitting, each UPS1 part was matched to a yeast part to create 10 sub-runs. Each of these sub-runs was used as input to a database search, as previously described, with each database search using a different decoy database.

Following the database search, and designating UPS1 peptides as relevant and yeast peptides as irrelevant, we applied to the resulting set of PSMs the selection procedures of subset-search, all-sub, group-FDR, and FSNS at a 5% FDR threshold.

The FDP in the FDR controlled set of PSMs was estimated by dividing the number of demonstrably incorrect PSMs (i.e., the number of times a yeast spectrum matched to a UPS1 peptide—additional detail can be found below) by the total number of discoveries. The FDP values from the ten sub-runs were used as input to the one-sided t-test. This process was repeated for the other two UPS1/yeast runs, yielding 30 sub-runs and three p-values.

## 3 Results

### 3.1 All-sub can fail to control FDR when the subset of interest is small

First we investigated whether subset-search, all-sub, and group-FDR each properly control FDR when the subset of interest is small. Note that we only test three methods because search-then-select has been previously shown to improperly control the FDR.^5–8^

To test these methods, we first estimated the FDP within an FDR-controlled set of PSMs. To do so, we computationally mixed together an irrelevant yeast run with a relevant UPS1 run. As a result, any yeast spectrum that is matched with a UPS1 peptide is demonstrably incorrect. This allows us to give a lower bound on the FDP. It should be noted that in this real data we do not precisely know the FDP; however, we designed our experiment so that in practice we believe our lower bound (i.e., estimated FDP) is fairly close to the actual, unknown one. Specifically, due to the large difference in size between the irrelevant yeast and relevant UPS1 database, we expect most incorrect PSMs that occur by chance would involve yeast peptides. After the FDP has been estimated, we performed a t-test to determine whether the mean of the FDP is significantly larger than the FDR threshold α. A mean FDP that is significantly larger than the FDR indicates that the corresponding FDR control method is probably invalid.

Our analysis suggests that all-sub fails to properly control the FDR (Table 5). Using the all-sub method, we calculated p-values of 0.0012944, 0.0008953, and 0.0024908 for the three different concatenated UPS1/yeast datasets. These p-values suggest that all-sub improperly controls the FDR because they are smaller than the Bonferroni corrected threshold of 0.004, where the uncorrected p-value threshold is 0.05 and n =12. Note that the same argument cannot be used against the remaining methods as their corresponding p-values are all above the Bonferroni corrected threshold. This of course does not prove that these three methods correctly control the FDR; however, in the absence of neighbors one can argue that all three of these methods satisfy the conditions that guarantee FDR control by TDC.^25^

**Table 5:** Assessing FDR control. Each p-value comes from a single t-test measuring whether the mean of the estimated FDP over the 10 sub-runs is significantly larger than the selected 5% FDR threshold. Each column uses a different computationally concatenated UPS1 and yeast run, and each row refers to a different FDR estimation procedure. Boldface values are significant at a Bonferroni corrected threshold of 0.004 (0.05/12). The analysis suggests that all-sub fails to control FDR.

| FDR control method | p-value 1 | p-value 2 | p-value 3 |
| --- | --- | --- | --- |
| subset-search | 0.25695 | 0.02642 | 0.20952 |
| all-sub | **0.0012944** | **0.0008953** | **0.0024908** |
| group-FDR | 0.05458 | 0.04435 | 0.05003 |
| filter then subset-neighbor search | 0.06294 | 0.02247 | 0.08400 |

### 3.2 Subset-search can fail to control the FDR in the presence of neighbors

Although our previous analysis indicates that subset-search properly handles a small subset of interest, we found that it can struggle to control the FDR in the presence of neighbor peptides. The issue is that, since subset-search does not search the spectra against the irrelevant peptides, a spectrum generated by an irrelevant neighbor peptide will likely receive a high score against the corresponding relevant peptide. As a result, these incorrect PSMs are more likely to be accepted as correct PSMs by target-decoy competition.

To give a concrete example, consider the theoretical MS2 spectrum of relevant ricin peptide “VGLPINQR” and irrelevant castor plant peptide “RIPLANGR” (Figure 2). These two peptides have a mass difference of approximately 12 ppm and have 69.56% of MS2 peaks in common. The PSM between the experimental scan (top row Figure 2) and the relevant peptide yields a combined p-value score of 1.25 × 10^−4^, which is larger (i.e., worse) than the combined p-value score, 1.13 × 10^−4^, of the PSM between the scan and the neighbor peptide. Hence, if the target database does not contain the neighbor peptides, then the database search would match this experimental scan with the relevant peptide even though it has an even better match with a neighbor peptide.

**Figure 2:**
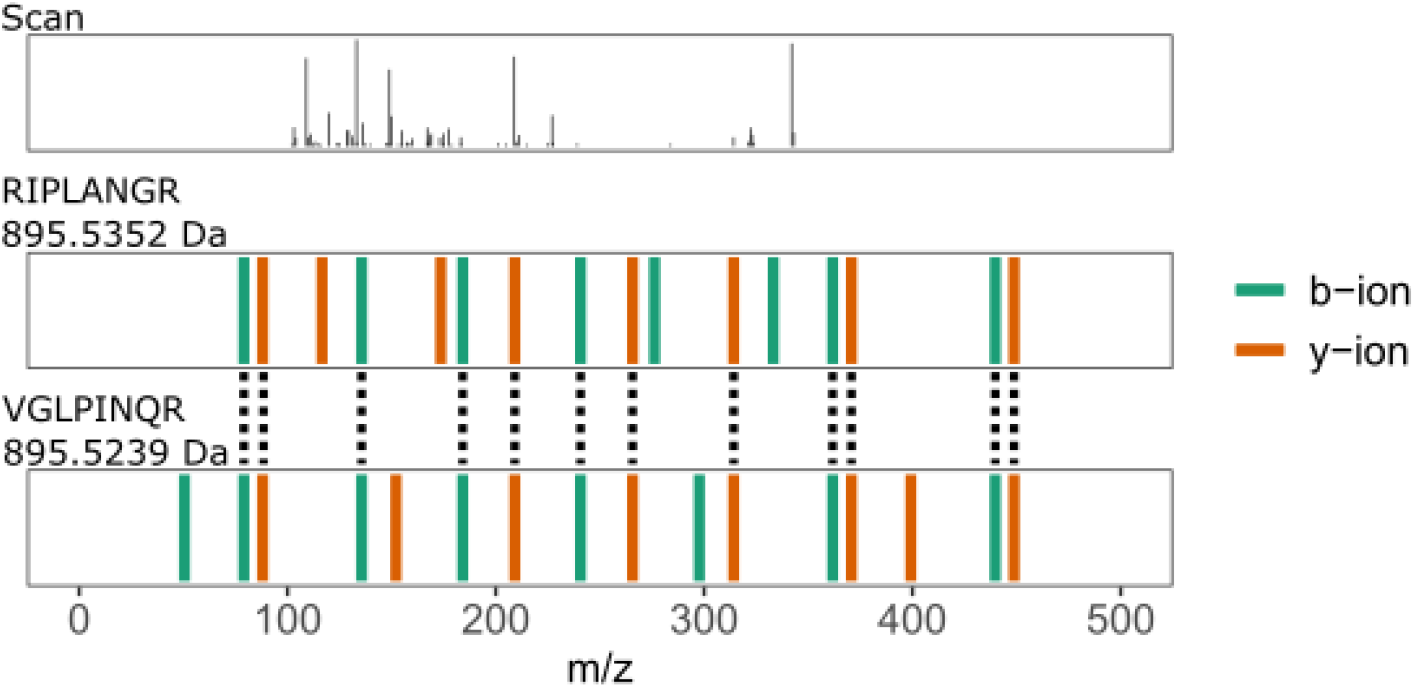
Example of a neighbor peptide. This figure plots an experimental spectrum with a precursor charge of two (top) along with the best scoring neighbor peptide (middle) and the best scoring relevant peptide (bottom). Peptide “VGLPINQR” is a relevant ricin peptide (Uniprot ID: B9T8T0) and peptide “RIPLANGR” is an irrelevant castor plant peptide (Uniprot ID: B9T289). The mass difference between the two peptides is approximately 12 ppm with the mass of “VGLPINQR” being 895.5239 Da and the mass of “RIPLANGR” being 895.5352 Da. These two peptides have ~70% of their MS2 peaks in common. Dotted lines connect MS2 peaks in the same 0.05 Da bin. The combined p-value score (lower is better) between the experimental scan with the relevant peptide is 1.25 × 10^−4^, whereas the combined p-value score between the scan and the neighbor peptide is 1.13 × 10^−4^.

To test our hypothesis that subset-search may be problematic in the presence of peptide neighbors, we investigated the confident set of PSMs, in each run, identified by subset-search at 1% FDR. For each such scan we asked whether that scan would have scored more highly if neighbor peptides had been present in the database. We repeated this process for each of the 54 different castor seed runs. We discovered that anywhere from 5 to 78 scans, in each set of confidently detected PSMs, switched their best scoring target from a relevant peptide to a neighbor peptide (Figure 3A). These scans comprise anywhere from 4.1% to 15.5% of the number of confidently identified scans (Figure 3B). Since the proportion of incorrect PSMs, due to lack of neighbor peptides in the database, is much larger than the FDR threshold of 1% FDR, this experiment suggests that subset fails to control the FDR in this scenario.

**Figure 3:**
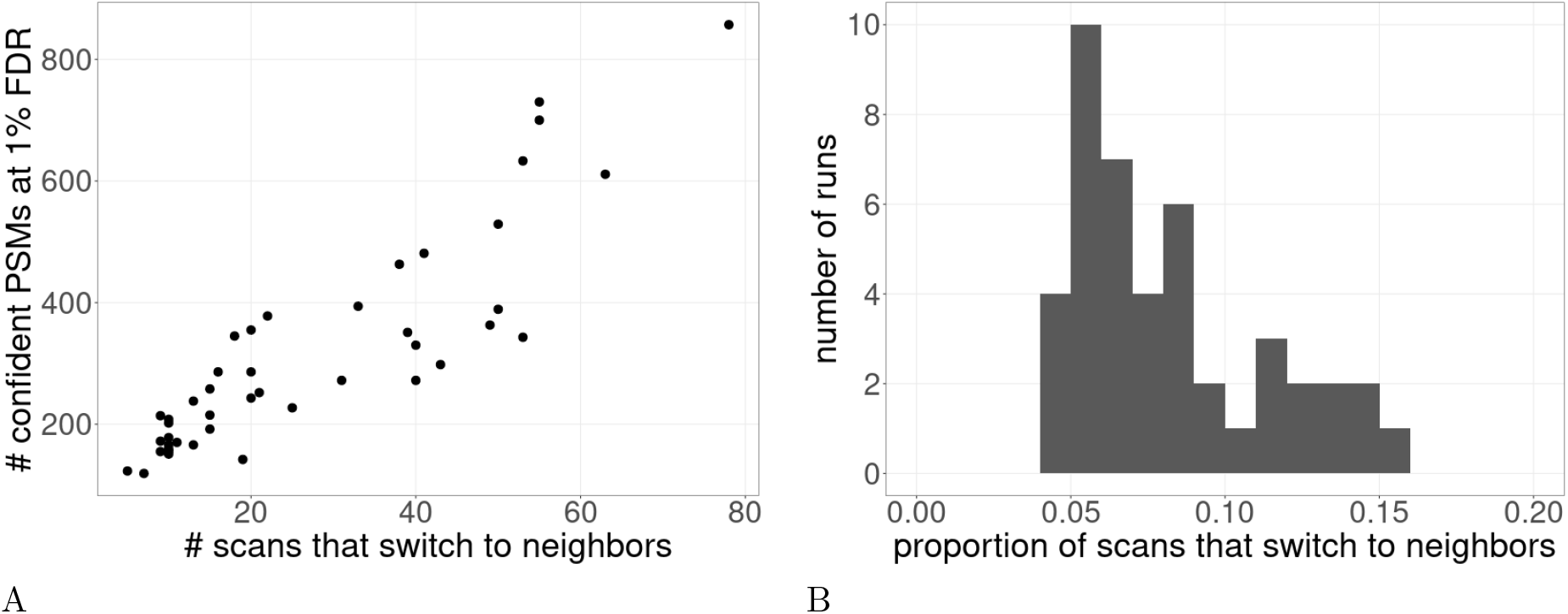
Magnitude of the neighbor peptide problem. This plot shows how subset-search does not properly control the FDR in the presence of many neighbors. We took the set of confidently detected PSMs, as detected by subset-search at 1% FDR. From this set of confident PSMs we determined the number of scans that would have scored better to a neighbor peptide if that peptide had been present in the database. This process was repeated for 54 different castor runs. (A) Each point is the number of scans in a run that would have matched to a neighbor peptide if neighbor peptides were searched as a function of the number of confident PSMs. (B) A histogram of the values from (A), where the x-axis is divided by the y-axis to obtain the proportion of scans that switch to neighbors. Note there are only 44 points because 10 runs had zero confident PSMs at 1% FDR.

### 3.3 FSNS controls for neighbor peptides and outperforms group-FDR

Our analysis so far suggests that, among existing methods, group-FDR is the only method that properly controls the FDR when the relevant database is much smaller than the irrelevant database. However, group-FDR suffers from low statistical power due to a high multiple testing hypothesis burden. This is because, in group-FDR, relevant spectra are searched against the entire database, which includes a large set of irrelevant peptides.^10^

For example, consider the ricin dataset. Looking at the difference in the median^2^ number of PSMs detected by subset-search and group-FDR at various FDR thresholds between 0–10% suggests that subset-search outperforms group-FDR across the entire q-value range of 0-10% (Figure 4). This observation motivated us to develop a new FDR control method that would share much of the power advantage of subset-search while correctly controlling the FDR by explicitly accounting for neighbors. Our method, called “filter then neighbor-subset search” (FSNS), consists of two separate database searches: a filtering and a primary search. In the filtering search, spectra are searched against relevant, neighbor, and irrelevant peptides. Following the search, PSMs that involve irrelevant targets or irrelevant decoys are extracted and FDR control via TDC is applied to this set of PSMs. Spectra that confidently match to a target irrelevant peptide are removed from consideration. In the subsequent primary search, the remaining spectra are searched against relevant and neighbor peptides. PSMs that involve relevant peptides or their associated decoys are extracted, and FDR control via TDC is applied to this set of PSMs to determine the list of confidently detected relevant PSMs. FSNS should in general offer more discoveries than group-FDR because spectra that match, in the filtering search, to either an irrelevant decoy or insignificantly to a relevant target are given a second chance to match a relevant or neighbor peptide in the primary search.

**Figure 4:**
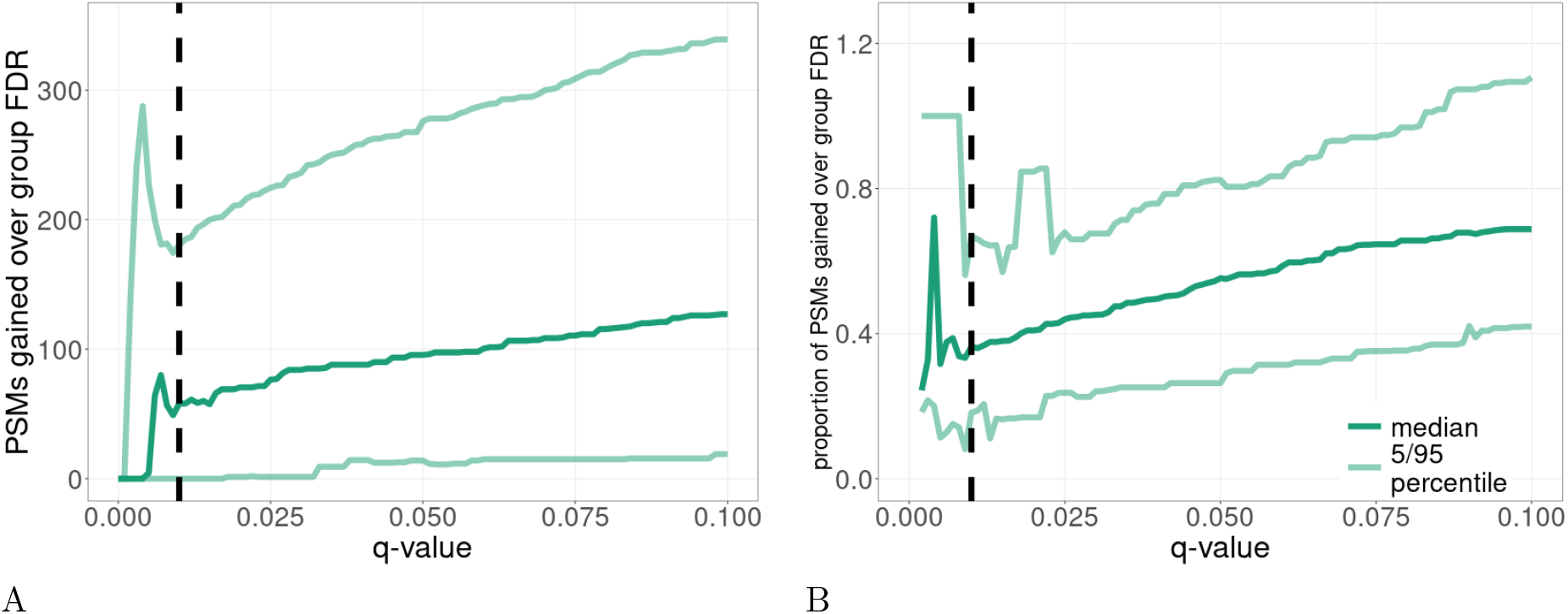
Comparison of subset-search and group-FDR. This figure compares the performance of subset-search against group-FDR in the ricin dataset with respect to (A) the number of PSMs and (B) the proportional increase in number of PSMs. For each mass spectrometry run, we determine the difference in the number of PSMs detected between subset-search and group-FDR at various FDR thresholds. After collating these values across all runs, we plot the median value and 5/95 percentiles over 54 runs. The vertical dashed line is at the conventional 1% FDR threshold. Note that the plotted values in B are undefined for some q-values near 0 where neither subset-search nor group-FDR detects any PSMs.

To test this in practice, we compared the performance of FSNS and group-FDR on three different datasets: ricin, non-concatenated yeast/UPS1, and human. These three datasets provide several examples of cases where the relevant portion of the database is a tiny portion of the overall database. The ricin example provides a real world example of a situation where an investigator is interested in a tiny subset of the possible database (where the relevant subset is on the order of 10^−5^ the size of the overall database). We use the yeast/UPS1 dataset as an example where the relevant portion is larger but still a small subset of the overall database (on the order of 10^−3^). Finally, we use the five iterations of the human database to show additional evidence (on the order of 10^−4^-10^−5^).

Empirically, we found that FSNS indeed generally outperforms group-FDR across the three datasets (Figure 5). In the ricin dataset, FSNS outperforms group-FDR across the entire FDR range of 0—10%. At a 1% FDR threshold, FSNS outperforms group-FDR by a median difference of 26.5 PSMs (Figure 5A). Looking at percent differences, FSNS outperforms group-FDR by a median percent difference of 16.2% (Figure 5C). On the other hand, we see slightly different results when we compare the performance of FSNS and group-FDR in the yeast/UPS1 dataset (Figure 5B and E). These two methods have comparable performance for small q-values (≤ 2.5%). However, for larger q-values (≥ 2.5%), FSNS outperforms group-FDR. Specifically, looking at a 5% q-value threshold, FSNS outperforms outperforms group-FDR by a median of 26.5 PSMs (3.1%). Finally, the five different iterations of the human dataset generally follow the trends previously described (Figure 5D and F). At low q-values, FSNS and group-FDR have similar power. Specifically at 1% FDR, FSNS and group-FDR have similar performance in four out of the five iterations. As the q-value threshold increases, FSNS steadily outperforms group-FDR. At a 5% q-value threshold FSNS improves upon group-FDR by 8 (3.55%), 10 (11.57%), 9 (7.73%), 7 (4.15%), and 12 PSMs (15.24%).

**Figure 5:**
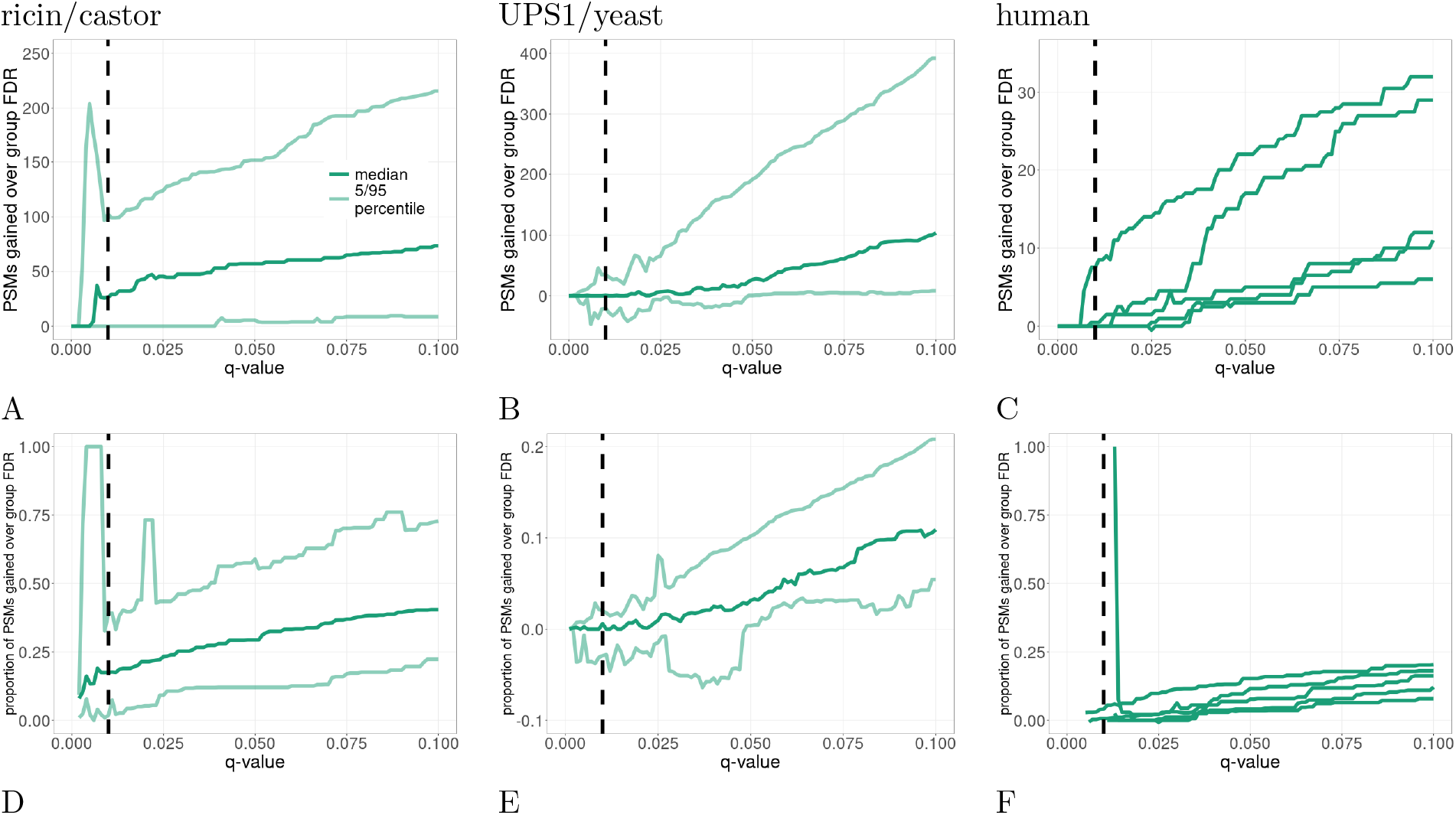
Comparison of FSNS and group-FDR. This plot compares the relative performance of FSNS against group-FDR for the ricin (A and D), UPS1/yeast (B and E), and human dataset (C and F). For each mass spectrometry run, we determine the difference in the number of PSMs (A—C) detected between FSNS and group-FDR at various q-value thresholds. In addition, we calculate the corresponding proportional increase (D—F). For D—F, we assigned a value of 1 when FSNS detects any number of PSMs while group-FDR detects O PSMs. The vertical dashed line is at the conventional 1% FDR threshold. Note the plotted values in D—F are undefined for some q-values near O, where neither FSNS nor group-FDR detects any PSMs. For the human data (C and F), we only plot the median lines, where each line represents a different relevant protein.

## 4 Discussion

When analyzing selection, or equivalently, FDR-controlling procedures, we should first evaluate their ability to control the FDR. We can then compare the power, or number of discoveries, among those procedures that indeed control the FDR.

It was already shown that search-then-select fails to control the FDR, and our analysis suggests that all-sub also fails to properly control the FDR when the relevant subset of peptides is small. Although a small relevant set of peptides is not a problem for subset-search on its own, we show that it does become so when the dataset contains a significant number of spectra that were generated by neighbor peptides.

Conceptually, the most natural source of neighbors peptides is from homologous sequences within and between proteomes. However, neighbor peptides are a more insidious problem than homology because two peptides with seemingly very different sequences can have very similar MS2 spectra. For example, although the peptide sequences in Figure 2 look different from each other, they have remarkably similar precursor masses and theoretical fragmentation spectra. Thus, neighbor peptides are a potential problem even in the absence of homology. In light of this observation, we feel it is too risky to recommend using subset-search hoping that the neighbors pose an insignificant problem.

This reasoning leaves us with group-FDR as the only established tool that can control the FDR in our context. That said, group-FDR (like search-then-select and all-sub) sacrifices power by searching all the spectra, including the ones generated by the relevant subset of peptides, against a database that includes a large number of irrelevant peptides. This loss of power was the motivation for introducing subset-search to begin with, and so our task here was to avoid most of this power loss while addressing the neighbor problem.

A simple fix to subset-search’s neighbor problem would have been to add the neighbors as an additional group (essentially, FSNS with no filtering or, equivalently, with *α′* = 0). The problem with this approach is that it still leaves us exposed to the problem of potential power loss in the presence of a very large number of spectra generated by irrelevant peptides (e.g., when looking for a particular post-translational modification present in a sample). Under such a scenario, due to the sheer number of such “irrelevant spectra” it is likely that a few spectra will generate some high scoring *random* PSMs when searched against the relevant peptides and their neighbors. Being random, about half of these PSMs will fall on the decoy side of the database, making it significantly more difficult for the few correct PSMs to stand out in the target-decoy competition analysis, especially if we use a poorly calibrated database search score function. Hence the idea of filtering: reduce the possibly significant chunk of the dataset that is irrelevant to us and that essentially just obfuscates the relevant analysis.

The detrimental effect of neighbor peptides on FDR estimation was previously discussed in the context of modifications.30 Here we demonstrated and addressed this effect in the context of subset search. We believe that the existence of neighbor peptides is also likely to have a large effect on the results of multi-pass searches.^26–29^ In these procedures, spectra are searched against a series of peptide databases. Such a setup can be problematic because peptides present in different databases can be neighbors of each other. As a result, spectra could incorrectly match well with a peptide found in the initial database search when in fact the best match would be found in subsequent searches. Future work needs to be done to quantify and address the neighbor peptides effect in the context of multi-pass search.

## 5 Acknowledgments

We thank Lindsay Pino for providing the UPS1 and yeast samples used in the FDR validity experiment. In addition we thank the University of Washington’s Proteomics Resource (UWPR95794) for instrument access. This work was funded by NIH award R01 GM121818.

## 6 Supporting Information

- **Supplemental File S1:** Pseudocode for the five different search procedures, plus three supplemental tables.
- **Supplemental File S2:** Python script (filterPSMsByGroup.py) used to select PSMs for subset search, group FDR, all-sub, and FSNS for input into target-decoy compeition and subsequent FDR estimation.
- **Supplemental File S3:** Python script (allSub.py) used to estimate all-sub q-values.

---

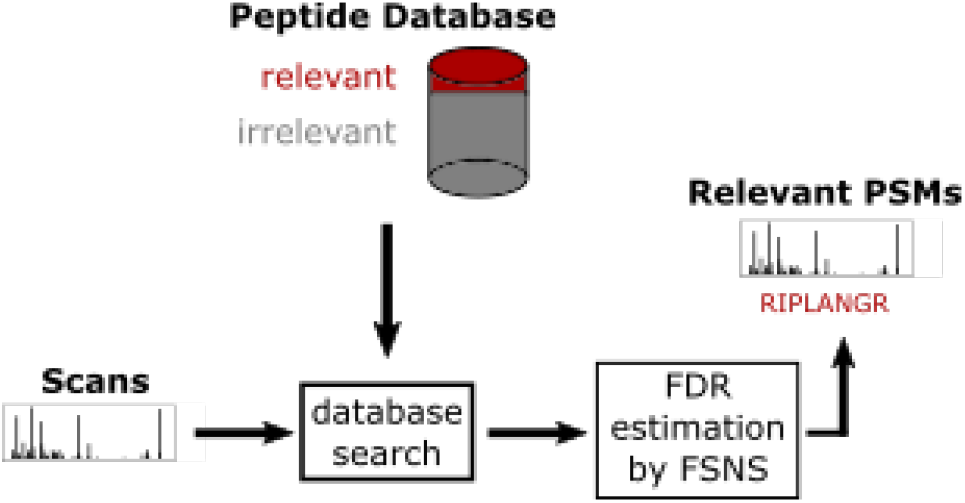

## Footnotes

1 Submission of the data is in progress.

2 We chose to use the median instead of the mean because we expect a different concentration of the relevant protein in each run, thereby changing the expected number of relevant PSMs.

## References

[1] J. E. Elias and S. P. Gygi. Target-decoy search strategy for increased confidence in large-scale protein identifications by mass spectrometry. Nature Methods, 4(3):207–214, 2007.

[2] J. E. Elias and S. P. Gygi. Target-decoy search strategy for mass spectrometry-based proteomics. Methods in Molecular Biology, 604(55-71), 2010.

[3] A. Z. Tremp, S. Saeed, V. Sharma, E. Lasonder, and J. T. Dessens. Plasmodium berghei laps form an extended protein complex that facilitates crystalloid targeting and biogenesis. Journal of Proteomics, 227:103925, 2020.

[4] S. E. Lindner, K. E. Swearingen, M. J. Shears, M. P. Walker, E. N. Vrana, K. J. Hart, A. M. Minns, P. Sinnis, R. L. Moritz, and S. H. I. Kappe. Transcriptomics and proteomics reveal two waves of translational repression during the maturation of malaria parasite sporozoites. Nature communications, 10(1):4964–4964, Oct 2019.

[5] X. Yi, F. Gong, and Y. Fu. Transfer posterior error probability estimation for peptide identification. BMC Bioinformatics, 21, May 2020.

[6] B. Efron. Simultaneous inference: When should hypothesis testing problems be combined? The Annals of Applied Statistics, pages 197–223, 2008.

[7] Y. Fu. Bayesian false discovery ratesfor post-translational modification proteomics. Statistics and Its Interface, 5:47–59, 2012.

[8] Y. Fu and X. Qian. Transferred subgroup false discovery rate for rare post-translational modifications detected by mass spectrometry. Molecular and Cellular Proteomics, 13(5):1359–1368, 2014.

[9] W. S. Noble. Mass spectrometrists should only search for peptides they care about. Nature Methods, 12(7):605–608, 2015.

[10] W. S. Noble and U. Keich. Response to “Mass spectrometrists should search for all peptides, but assess only the ones they care about”. Nature Methods, 14(7):644, 2017.

[11] B. Efron. Microarrays, empirical bayes and the two-groups model. Statistical Science, 23(1):1–22, 2008.

[12] A. Sticker, L. Martens, and L. Clement. Mass spectrometrists should search for all peptides, but assess only the ones they care about. Nature Methods, 14(7):643–644, 2017.

[13] X. Yi, B. Wang, Z. An, F. Gong, J. Li, and Y. Fu. Quality control of single amino acid variations detected by tandem mass spectrometry. Journal of Proteomics, 187:144–151, 2018.

[14] C. Ramus, A. Hovasse, M. Marcellin, A. Hesse, E. Mouton-Barbosa, D. Bouyssia, S. Vaca, C. Carapito, K. Chaoui, C. Bruley, J. Garin, S. Cianfaani, M. Ferro, A. V. Dorssaeler, O. Burlet-Schiltz, C. Schaeffer, Y. Coutaa, and A. Gonzalez de Peredo. Spiked proteomic standard dataset for testing label-free quantitative software and statistical methods. Data in Brief, 6, March 2016.

[15] E. D. Merkley, S. C. Jenson, J. S. Arce, A. M. Melville, O. P. Leiser, D. S. Wunschel, and K. L. Wahl. Ricin-like proteins from the castor plant do not influence liquid chromatography-mass spectrometry detection of ricin in forensically relevant samples. Toxicon, 140:18–31, 2017.

[16] J. Doellinger, A. Schneider, M. Hoeller, and P. Lasch. Sample preparation by easy extraction and digestion (speed) - a universal, rapid, and detergent-free protocol for proteomics based on acid extraction. Molecular & Cellular Proteomics, 19(1):209–222, 2020.

[17] J. R. Wisniewski, Alexandre Zougman, Nagarjuna Nagaraj, and Matthias Mann. Universal sample preparation method for proteome analysis. Nature Methods, 13:359–362, 2009.

[18] M. Sielaff, J. Kuharev, T. Bohn, J. Hahlbrock, T. Bopp, S. Tenzer, and U. Distler. Evaluation of fasp, sp3, and ist protocols for proteomic sample preparation in the low microgram range. Journal of Proteome Research, 16(11):4060–4072, 2017.

[19] W. H. McDonald, D. L. Tabb, R. G. Sadygov, M. J. MacCoss, J. Venable, J. Graumann, J. R Johnson, D. Cociorva, and J. R. Yates, III. MS1, MS2, and SQT-three unified, compact, and easily parsed file formats for the storage of shotgun proteomic spectra and identifications. Rapid Communications in Mass Spectrometry, 18:2162–2168, 2004.

[20] D. Kessner, M. Chambers, R. Burke, D. Agnus, and P. Mallick. Proteowizard: open source software for rapid proteomics tools development. Bioinformatics, 24(21):2534–2536, 2008.

[21] S. McIlwain, K. Tamura, A. Kertesz-Farkas, C. E. Grant, B. Diament, B. Frewen, J. J. Howbert, M. R. Hoopmann, L. Kall, J. K. Eng, M. J. MacCoss, and W. S. Noble. Crux: rapid open source protein tandem mass spectrometry analysis. Journal of Proteome Research, 13(10):4488–4491, 2014.

[22] C. Y. Park, A. A. Klammer, L. Käll, M. P. MacCoss, and W. S. Noble. Rapid and accurate peptide identification from tandem mass spectra. Journal of Proteome Research, 7(7):3022–3027, 2008.

[23] A. Lin, J. J. Howbert, and W. S. Noble. Combining high-resolution and exact calibration to boost statistical power: A well-calibrated score function for high-resolution ms2 data. Journal of Proteome Research, 17:3644–3656, 2018.

[24] D. H. May, K. Tamura, and W. S. Noble. Param-Medic: A tool for improving MS/MS database search yield by optimizing parameter settings. Journal of Proteome Research, 16(4):1817–1824, 2017.

[25] K. He, Y. Fu, W.-F. Zeng, L. Luo, H. Chi, C. Liu, L.-Y. Qing, R.-X. Sun, and S.-M. He. A theoretical foundation of the target-decoy search strategy for false discovery rate control in proteomics. arXiv, 2015.

[26] A. Kertesz-Farkas, U. Keich, and W. S. Noble. Tandem mass spectrum identification via cascaded search. Journal of Proteome Research, 14(8):3027–3038, 2015.

[27] P. Kumar, J. E. Johnson, C. Easterly, S. Mehta, R. Sajulga, B. Nunn, P. D. Jagtap, and T. J. Griffin. A sectioning and database enrichment approach for improved peptide spectrum matching in large, genome-guided protein sequence databases. Journal of Proteome Research, 19(7):2772–2785, 2020. PMID: 32396365.

[28] S. Woo, S. W. Cha, S. Bonissone, S. Na, D. L. Tabb, P. A. Pevzner, and V. Bafna. Advanced proteogenomic analysis reveals multiple peptide mutations and complex immunoglobulin peptides in colon cancer. Journal of Proteome Research, 14(9):3555–3567, 2015. PMID: 26139413.

[29] P. Jagtap, J. Goslinga, J. A. Kooren, T. McGowan, M. S. Wroblewski, S. L. Seymour, and T. J. Griffin. A two-step database search method improves sensitivity in peptide sequence matches for metaproteomics and proteogenomics studies. Proteomics, 13(8):1352–1357, 2013.

[30] A. T. Kong, F. V. Leprevost, D. M. Avtonomov, D. Mellacheruvu, and A. I. Nesvizhskii. MSFragger: ultrafast and comprehensive peptide identification in mass spectrometry-based proteomics. Nature Methods, 14(5):513–520, 2017.

