## Supplemental File 1 for "Improving power while controlling the false discovery rate when only a subset of peptides are relevant"

**Algorithm 1 Search-then-select**

**procedure** SEARCHTHENSELECT$(S, \mathcal{T}, \mathcal{D}, \mathcal{T}_r, \alpha)$
    $(M, P) := \text{SEARCH}(S, \mathcal{T} \cup \mathcal{D})$
    $A := \text{CONTROLFDR}(M, P, \alpha)$
    $R := \{(s_i, p_i, m_i) \mid a_i = 1, p_i \in \mathcal{T}_r\}$
    **return** R
**end procedure**

**Algorithm 2 Subset-search**

**procedure** SUBSETSEARCH$(S, \mathcal{T}_r, \mathcal{D}_r, \alpha)$
    $(M, P) := \text{SEARCH}(S, \mathcal{T}_r \cup \mathcal{D}_r)$
    $A := \text{CONTROLFDR}(M, \alpha)$
    $R := \{(s_i, p_i, m_i) \mid a_i = 1\}$
    **return** R
**end procedure**

**Algorithm 3 Group-FDR**

**procedure** GROUPFDR $(S, \mathcal{T}_r, \mathcal{D}_r, \mathcal{T}_i, \mathcal{D}_i, \alpha)$
    $(M, P) := \text{SEARCH}(S, \mathcal{T}_r \cup \mathcal{T}_i \cup \mathcal{D}_r \cup \mathcal{D}_i)$
    $(M^1, P^1) := \{(m_i, p_i) \mid p_i \in \mathcal{T}_r \cup \mathcal{D}_r\}$
    $A := \text{CONTROLFDR}(M^1, \alpha)$
    $R := \{(s_i^1, p_i^1, m_i^1) \mid a_i = 1\}$
    **return** R
**end procedure**

**Algorithm 4 All-sub**

**procedure** ALLSUB($S, \mathcal{T}, \mathcal{D}, \mathcal{T}_r, \alpha$)
  $(M, P) := \text{SEARCH}(S, \mathcal{T} \cup \mathcal{D})$
  $A := \text{CONTROLFDR2}(M, \alpha)$
  $R := \{(s_i, p_i, m_i) \mid a_i = 1\}$
  **return** R
**end procedure**

**Algorithm 5 Filter then subset-neighbor search**

**procedure** FILTERSUBSETNEIGHBORSEARCH($S, \mathcal{T}_r, \mathcal{D}_r, \mathcal{T}_i, \mathcal{D}_i, \mathcal{T}_n, \mathcal{D}_n, \alpha', \alpha$)
  $(M, P) := \text{SEARCH}(S, \mathcal{T}_r \cup \mathcal{T}_i \cup \mathcal{T}_n \cup \mathcal{D}_r \cup \mathcal{D}_i \cup \mathcal{D}_n)$
  $(M^1, P^1) := \{(m_i, p_i) \mid p_i \in \mathcal{T}_i \cup \mathcal{D}_i\}$
  $A := \text{CONTROLFDR}(M^1, \alpha')$
  $S^1 := \{s_i \mid a_i = 0\}$
  $(M^2, P^2) := \text{SEARCH}(S^1, \mathcal{T}_r \cup \mathcal{T}_n \cup \mathcal{D}_r \cup \mathcal{D}_n)$
  $(M^3, P^3) := \{(m_i, p_i) \mid p_i \in \mathcal{T}_r \cup \mathcal{D}_r\}$
  $A^1 := \text{CONTROLFDR}(M^3, \alpha)$
  $R := \{(s_i^1, p_i^1, m_i^1) \mid a_i^1 = 1\}$
  **return** R
**end procedure**

| file name | # scans |
|---|---|
| UPS1_12500amol_R1.ms2 | 39718 |
| UPS1_12500amol_R2.ms2 | 39682 |
| UPS1_12500amol_R3.ms2 | 39782 |
| UPS1_25000amol_R1.ms2 | 40856 |
| UPS1_25000amol_R2.ms2 | 40512 |
| UPS1_25000amol_R3.ms2 | 40439 |
| UPS1_50000amol_R1.ms2 | 41833 |
| UPS1_50000amol_R2.ms2 | 41653 |
| UPS1_50000amol_R3.ms2 | 41665 |
| UPS1_5000amol_R1.ms2 | 37918 |
| UPS1_5000amol_R2.ms2 | 38046 |
| UPS1_5000amol_R3.ms2 | 38052 |

Table 1: **yeast/UPS1 data.** The number of scans found in each run. This is for the samples where UPS1 was spiked into yeast.

| *file name | cultivar | preparation method | # scans | *file name | cultivar | preparation method | # scans |
|---|---|---|---|---|---|---|---|
| Zanz_1_1_03Jun16 | *Zanzibarensis* | M0 | 15735 | GCH4_1_1_03Jun16 | GCH4 | M0 | 21634 |
| Zanz_1_2_03Jun16 | *Zanzibarensis* | M0 | 17138 | GCH4_1_2_03Jun16 | GCH4 | M0 | 16765 |
| Zanz_1_3_03Jun16 | *Zanzibarensis* | M0 | 18834 | GCH4_1_3_03Jun16 | GCH4 | M0 | 18663 |
| Zanz_2_1_03Jun16 | *Zanzibarensis* | M0 | 19760 | GCH4_2_1_03Jun16 | GCH4 | M0 | 19830 |
| Zanz_2_2_03Jun16 | *Zanzibarensis* | M0 | 18986 | GCH4_2_2_03Jun16 | GCH4 | M0 | 24944 |
| Zanz_2_3_03Jun16 | *Zanzibarensis* | M0 | 16222 | GCH4_2_3_03Jun16 | GCH4 | M0 | 21667 |
| Zanz_3_1_03Jun16 | *Zanzibarensis* | M0 | 18900 | GCH4_3_1_03Jun16 | GCH4 | M0 | 20267 |
| Zanz_3_2_03Jun16 | *Zanzibarensis* | M0 | 21368 | GCH4_3_2_03Jun16 | GCH4 | M0 | 22246 |
| Zanz_3_3_03Jun16 | *Zanzibarensis* | M0 | 20315 | GCH4_3_3_03Jun16 | GCH4 | M0 | 25330 |
| TMVCH1_1_1_03Jun16 | TMVCH1 | M0 | 14893 | 200_1_03Jun16 | 200 | M0 | 13742 |
| TMVCH1_1_2_03Jun16 | TMVCH1 | M0 | 20086 | 200_2_03Jun16 | 200 | M0 | 17822 |
| TMVCH1_1_3_03Jun16 | TMVCH1 | M0 | 23470 | 200_3_03Jun16 | 200 | M0 | 6158 |
| TMVCH1_2_1_03Jun16 | TMVCH1 | M0 | 19125 | 5952_1_03Jun16 | 592 | M0 | 14951 |
| TMVCH1_2_2_03Jun16 | TMVCH1 | M0 | 23804 | 592_2_03Jun16 | 592 | M0 | 16157 |
| TMVCH1_2_3_03Jun16 | TMVCH1 | M0 | 20689 | 592_3_03Jun16 | 592 | M0 | 14125 |
| TMVCH1_3_1_03Jun16 | TMVCH1 | M0 | 16456 | 611_1_03Jun16 | 611 | M0 | 10724 |
| TMVCH1_3_2_03Jun16 | TMVCH1 | M0 | 25938 | 611_2__03Jun16 | 611 | M0 | 13621 |
| TMVCH1_3_3_03Jun16 | TMVCH1 | M0 | 16414 | 611_3_03Jun16 | 611 | M0 | 8454 |
| 1_M0_AM_R1_7Mar16 | PNNL | M0 | 5076 | 829_1_03Jun16 | 829 | M0 | 17343 |
| 2_M2_JA_R3_7Mar16 | PNNL | M2 | 16784 | 3_M0_JA_R3_7Mar16 | 829 | M0 | 18592 |
| 3_M0_JA_R3_7Mar16 | PNNL | M0 | 15859 | 829_3_03Jun16 | 829 | M0 | 9475 |
| 3_M0_JA_R3_B_03Jun16 | PNNL | M0 | 22015 | 8_M1_JA_R2_7Mar16 | PNNL | M1 | 15669 |
| 4_M1_AM_R1_7Mar16 | PNNL | M1 | 13188 | 9_M4_AM_R1_7Mar16 | PNNL | M4 | 14892 |
| 5_M2_AM_R1_7Mar16 | PNNL | M2 | 15358 | 9_M4_JA_R2_7Mar16 | PNNL | M4 | 16140 |
| 6_M0_JA_R2_7Mar16 | PNNL | M0 | 16728 | 9_M4_JA_R3_7Mar16 | PNNL | M4 | 15472 |
| 6_M0_JA_R2_B_03Jun16 | PNNL | M0 | 21544 | 10_M2_JA_R2_7Mar16 | PNNL | M2 | 18358 |
| 7_M1_JA_R3_7Mar16 | PNNL | M1 | 17973 | 16_M3_03Jun16 | PNNL | M3 | 20583 |

Table 2: **Ricin data.** The number of scans found in each ricin run. *All file names start with "Rcom_" and end with either "_Samwise_16-03-32.ms2" or "_Samwise_15-08-55.ms2".

| file name | # scans |
|---|---|
| 217_2018_ZBS6_HeLa_ISD_1.ms2 | 89162 |
| 220_2018_ZBS6_HeLa_SPEED_1.ms2 | 87981 |
| 222_2018_ZBS6_HeLa_FASP_2.ms2 | 87398 |
| 223_2018_ZBS6_HeLa_ISD_2.ms2 | 86869 |
| 225_2018_ZBS6_HeLa_SP3_2.ms2 | 90238 |
| 226_2018_ZBS6_HeLa_SPEED_2.ms2 | 88086 |
| 228_2018_ZBS6_HeLa_FASP_3.ms2 | 87255 |
| 229_2018_ZBS6_HeLa_ISD_3.ms2 | 87835 |
| 231_2018_ZBS6_HeLa_SP3_3.ms2 | 90546 |
| 232_2018_ZBS6_HeLa_SPEED_3.ms2 | 87703 |
| 234_2018_ZBS6_HeLa_FASP_1.ms2 | 87797 |
| 235_2018_ZBS6_HeLa_SP3_1.ms2 | 89253 |

Table 3: **Human data.** The number of scans found in each run.
